# Modelling cell guidance and curvature control in evolving biological tissues

**DOI:** 10.1101/2020.07.10.197020

**Authors:** Solene G.D. Hegarty-Cremer, Matthew J. Simpson, Thomas L. Andersen, Pascal R. Buenzli

## Abstract

Tissue geometry is an important influence on the evolution of many biological tissues. The local curvature of an evolving tissue induces tissue crowding or spreading, which leads to differential tissue growth rates, and to changes in cellular tension, which can influence cell behaviour. Here, we investigate how directed cell motion interacts with curvature control in evolving biological tissues. Directed cell motion is involved in the generation of angled tissue growth and anisotropic tissue material properties, such as tissue fibre orientation. We develop a new cell-based mathematical model of tissue growth that includes both curvature control and cell guidance mechanisms to investigate their interplay. The model is based on conservation principles applied to the density of tissue synthesising cells at or near the tissue’s moving boundary. The resulting mathematical model is a partial differential equation for cell density on a moving boundary, which is solved numerically using a hybrid front-tracking method called the cell-based particle method. The inclusion of directed cell motion allows us to model new types of biological growth, where tangential cell motion is important for the evolution of the interface, or for the generation of anisotropic tissue properties. We illustrate such situations by applying the model to simulate both the resorption and infilling components of the bone remodelling process, and to simulate root hair growth. We also provide user-friendly MATLAB code to implement the algorithms.

## 1. Introduction

Understanding the mechanisms controlling the generation of biological tissue is an important challenge in biomechanics and mechanobiology (Ambrosi et al., 2019) with key applications in tissue engineering and developmental biology (O’Brien, 2011, Dzobo et al., 2018). Tissue geometry influences the generation of new tissue, particularly the rate of tissue growth and the organisation of tissue material (Curtis and Varde, 1964, Dunn and Heath, 1976, Kollmannsberger et al., 2011). Several tissue growth experiments show that the rate of tissue progression is strongly dependent on tissue curvature. These findings apply to bioscaffold pore infilling (Bidan et al., 2013, 2016, Guyot et al., 2014, Ripamonti and Roden, 2010), wound healing (Poujade et al., 2007, Rolli et al., 2012), tumour growth (Lowengrub et al., 2010), and bone remodelling (Martin, 2000, Alias and Buenzli, 2018). This proportionality of growth rate and curvature may be caused by the crowding and spreading of cells and tissue material due to spatial constraints, and curvature-dependent tissue surface tension influencing cell proliferation rates (Nelson et al., 2005, Rumpler et al., 2008, Haeger et al., 2015, Alias and Buenzli, 2017, Buenzli et al., 2020).

In addition to the collective influence of curvature on tissue progression, other factors such as mechanical or chemical cues in the environment as well as cell-scale geometrical features can induce individual cell responses including directed cell migration. Mechanical cues include viscoelasticity (Chaudhuri et al., 2016), surface stiffness (Pelham and Wang, 1997, Lo et al., 2000, Discher et al., 2005, Engler et al., 2006), or surface mechanical stretch (Trepat et al., 2007, Livne et al., 2014). Chemical cues include signalling molecules inducing attractive or repulsive chemical gradients (Haeger et al., 2015), and cell-scale geometrical cues include geometrical guidance such as curvotaxis (Callens et al., 2020) and surface roughness gradients (Martin et al., 1995, Deligianni et al., 2001). While the collective influence of curvature on tissue growth and the effects of environmental cues on cell guidance mechanisms are well studied in isolation, how these processes interact during the generation of new biological tissue remains poorly understood.

In this work we develop a new mathematical model which explicitly includes both the collective influence of curvature and directed cell guidance mechanisms. The addition of directed cell guidance allows us to model new types of biological growth, which cannot be generated by existing mathematical models where the tissue interface progresses in the normal direction only (Bidan et al., 2013, Guyot et al., 2014, Alias and Buenzli, 2017, Callens et al., 2020).

Indeed, the growth of several tissues involves directed cell motion where cells move tangentially along the tissue surface (Figure 1). For example, shells, horns, and tusks with a spiralling structure are generated by tissue being secreted at an angle to the base membrane (Figure 1a) (Skalak et al., 1982, 1997). Tangential cell velocity may also be responsible for the generation of anisotropies in tissue material properties by aligning tissue fibrils with respect to the cells motion (Figure 2). In lamellar bone, the so-called twisted plywood structure of collagen fibrils may be due to the osteoblasts (bone secreting cells) changing direction of motion during bone infilling (Martin et al., 2004) (Figure 1c). Finally, tangential cell motion is suspected to occur in bone resorption to keep osteoclasts at the front of the resorption cone (Figure 1c).

**Figure 1:**
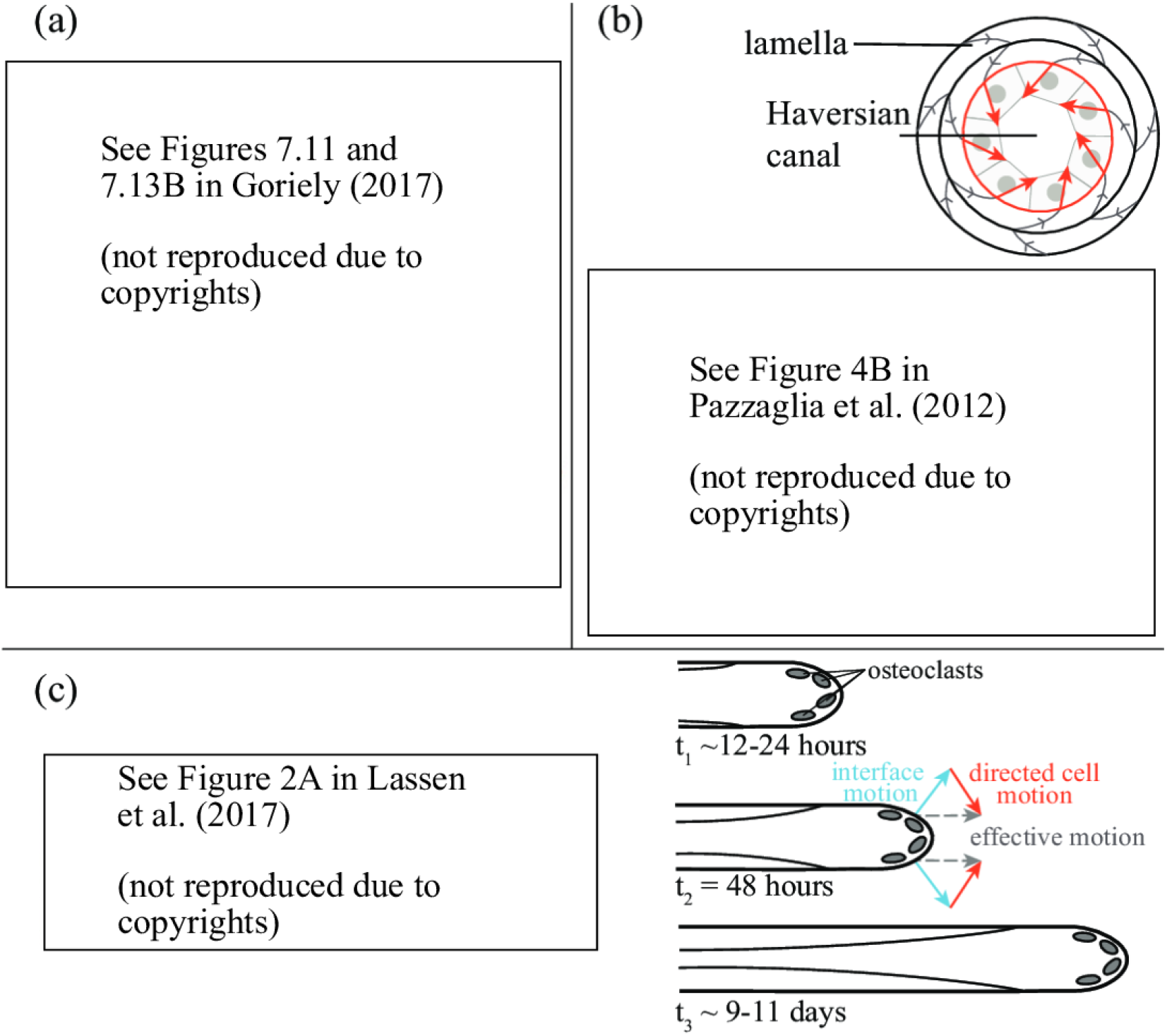
Tangential cell movement in tissue growth. (a) Shells grow by secretion of new tissue at their base (mantle) at an angle to create spiralling structures (reproduced with permissions from Goriely (2017)). (b) In lamellar bone, successive tissue layers possess different collagen fibril orientations which suggest changes in the tangential motion of osteoblasts during bone formation (reproduced with permissions from Pazzaglia et al. (2012) and Schrof et al. (2014)). (c) Resorption cavities during bone resorption maintain a stable resorption front shape at the tip. Since the dissolution process of bone by osteoclasts is expected to occur in the normal direction, this suggests osteoclasts are subject to cell guidance signals toward the cavity centerline. An example of the serial section of a cutting cone, immunostained (black) for an osteoclastic marker, obtained from Lassen et al. (2017) and schematic of an evolving Haversian system, after Jaworski and Hooper (1980).

**Figure 2:**
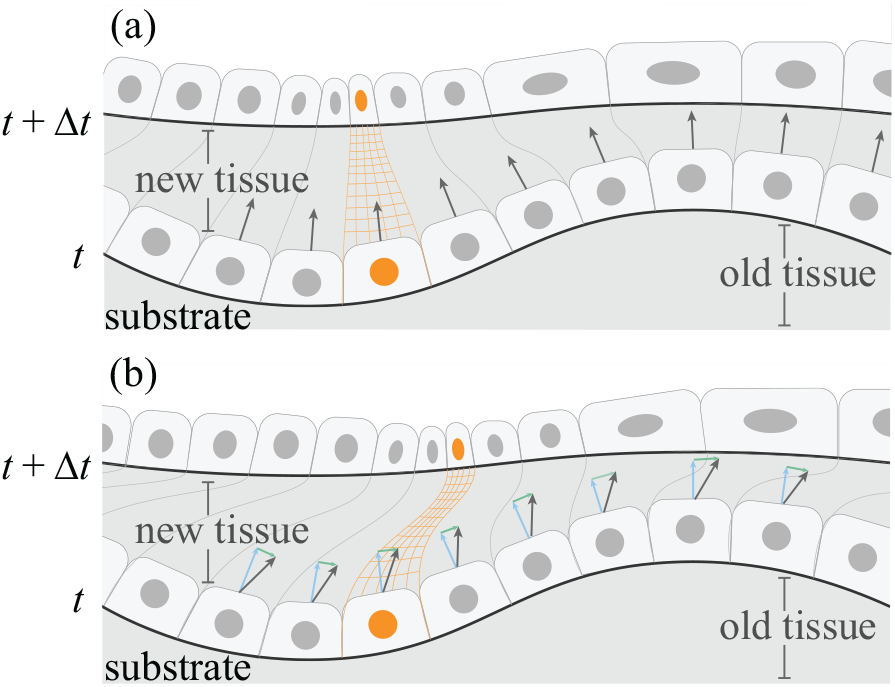
Schematic illustrating the crowding and spreading effect of curvature and the influence of tangential motion for tissue material properties; (a) shows only movement in the normal direction and the resulting changes in density; (b) includes both curvature control and cell guidance, meaning the cells crowd and spread and also undergo directed motion, creating anisotropies in tissue material properties (thin orange lines).

Mathematically, the evolution of *smooth* interfaces can be described by the normal velocity of the interface only (Sethian, 1999). However, biological tissue interfaces may develop cusps and sharp edges (Skalak et al., 1997, Alias and Buenzli, 2017, Goriely, 2017). When these move at an angle to their base, one is required to consider a more general tissue interface velocity that includes a tangential component to avoid the emergence of singularities in the governing equation for tissue growth velocity (Skalak et al., 1997).

Many existing models of geometric control of tissue growth consider the geometry of the tissue substrate only, so that cell guidance mechanisms and cell crowding effects are not modelled explicitly (Skalak et al., 1982, 1997, Rumpler et al., 2008, Bidan et al., 2012, 2013, Gamsjager, 2013, Guyot et al., 2014, Goriely, 2017, Ehrig et al., 2019). Here, we consider the cell-based mathematical model of Alias and Buenzli (2017), which explicitly accounts for curvature-induced cell crowding and spreading, and we generalise this model to allow for tangential cell motion. We derive the model from general conservation properties imposed on cells, which allows us to explicitly include cell behaviours. To our knowledge, no mathematical model currently includes both the effect of curvature on collective cell crowding and spreading and tangential cell motion mechanisms.

The model of Alias and Buenzli (2017) is also extended to three dimensions and the governing equations are derived in covariant form. The model derived is a partial differential equation (PDE) for the density of cells to be solved on a moving boundary, which represents the evolving tissue surface. This problem is numerically solved to explore several situations in which tangential cell guidance mechanisms are added. We demonstrate that with the addition of tangential cell advection, new biologically relevant tissue growth phenomena can be modelled, such as bone resorption, the generation of different fibre orientations in lamellar bone, and root hair growth.

## 2. Description of the model

Tissue growth usually occurs by cells synthesising new tissue close to the tissue’s interface. To determine general evolution equations for the density of tissue-synthesising cells subject to normal and tangential motion, we consider the case where the tissue-synthesising cells are attached to the tissue interface and described by a surface density, *ρ* (number of cells per unit surface). The motion of the interface transports the cells in space and the cells may additionally move laterally with respect to the material points of the surface. The motion of the interface is considered to be due to new tissue being synthesised in the wake of these surface-bound cells (Figure 2). This situation occurs for example in wound healing, bone remodelling, bioscaffold pore infilling, and tumour growth (Guyot et al., 2014, Bidan et al., 2013, Lowengrub et al., 2010, Poujade et al., 2007, Rumpler et al., 2008) where tissue-synthesing cells are located at or near the tissue interface. The normal velocity of the tissue interface, *u_n_***n** where **n** is the outward-facing unit surface normal, is given by

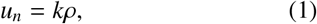

where *k* is the tissue-synthesising cells’ secretory rate (volume of new tissue synthesised per unit time per cell) (Buenzli, 2015). Tissue resorption can also be modelled by assuming *k* to be negative. Although in this derivation we take *ρ* to denote the density of tissue-secreting cells, other situations can be modelled if *ρ* is taken as the density of other surface-bound tissue-secreting entities, such as secretory vesicles (see Section 3.4).

During the evolution of the tissue, the interface may stretch locally depending on its curvature (Figure 2a), and this will induce changes in cell density. Convex areas of the tissue substrate result in cells spreading whereas concave areas of the tissue substrate result in cells crowding. In addition, cell guidance mechanisms superimpose lateral cell motion with respect to the tissue interface. Directional tissue growth may therefore result from a combination of interface motion and lateral cell motion (Figures 2b and 3).

**Figure 3:**
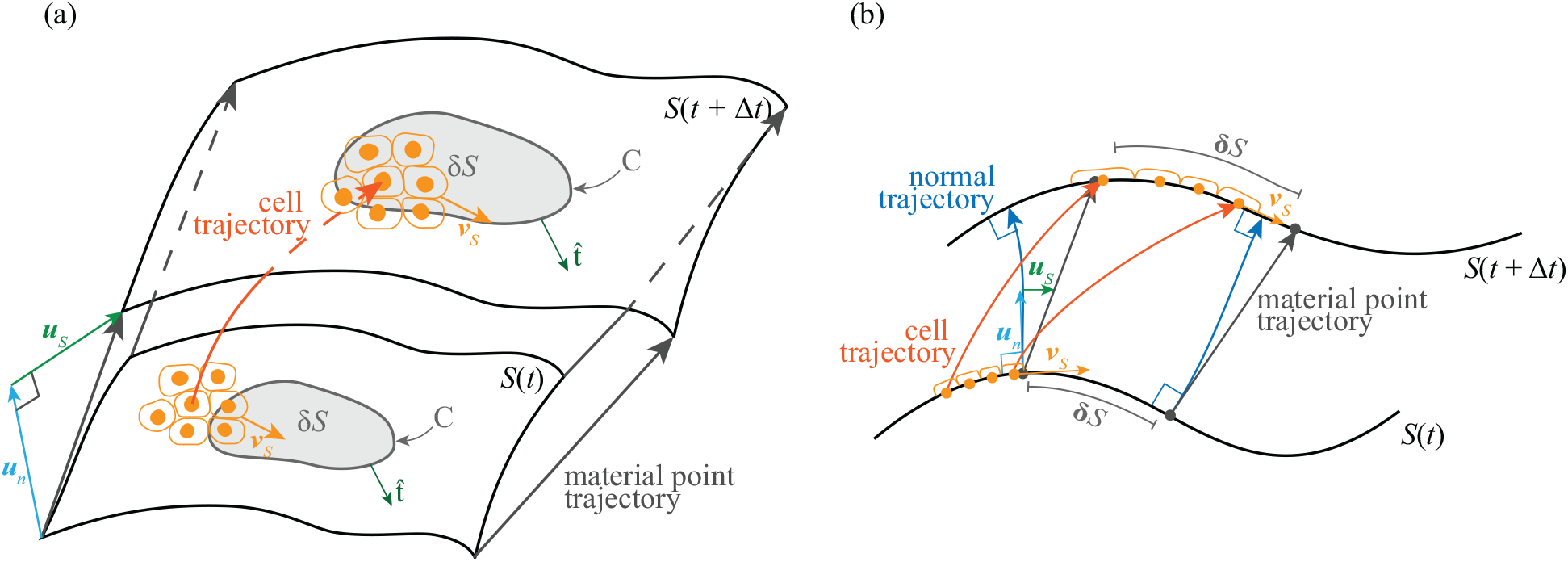
(a) Schematic of two dimensional surface portion being considered. The curve *C* surrounding δ*S* is illustrated as well as its outward facing normal 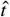. (b) One dimensional schematic of portion of interface being examined. The normal and tangential components of the surface velocity are annotated in blue and green respectively. Illustrative cells are included in orange, with the tangential flux of cells into δ**S** annotated in orange. The grey arrows indicate the material trajectories of the surface. Cell trajectories and normal trajectories are also annotated.

The tangential velocities of both the interface and the cells can be chosen to describe multiple biological tissue evolution scenarios. The tangential velocity of the cells, **v**_*S*_, can represent for example epithelial cells moving with respect to a basal membrane which may itself be transported in space with velocity **u**. Biological situations where cells are not physically transported by a moving tissue interface may be modelled by assuming that there is no tangential movement of the interface (**u**_*S*_ = 0, where **u**_*S*_ is the tangential component of **u**) while cells may still have tangential velocity (**v**_*S*_ ≠ 0). This can occur in the case of bone resorption for example, where material points of the bone interface do not move laterally but osteoclasts living on the interface may (Lassen et al., 2017). It is important to note that although the velocity of the tissue surface and the cells may not be distinguishable for modelling the evolution of the tissue interface and changes in cell density, the distinction between these velocities can be important for modelling the tissue material properties produced (Figure 2b, Buenzli (2016)), as we will illustrate in our application of the model to bone formation (see Section 3.2).

The tissue interface is denoted by *S* (*t*) and *ρ*(**r**_*S*_, *t*) denotes the surface density of the tissue-synthesising cells, at position **r**_*S*_ on *S* (*t*). We formally define *ρ*(**r**_*s*_, *t*) by considering an infinitesimal element of surface δ*S* at position **r**_*S*_ of *S* (*t*), and the number of cells living on this area, δ*N*. It is important to choose δ*S* small enough to capture heterogeneous densities but large enough to contain a sufficient number of cells to define a continuous surface density of cells, such that

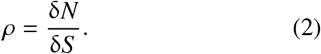

We now derive a conservation law for the surface density of cells living on the evolving surface as the tissue evolves. To do so, we consider the material derivative of *ρ* following the material trajectories, **r**_*S*_ (*t*), of the surface *S* (*t*), defined as

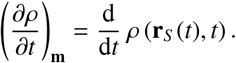

The material derivative obeys standard rules of differentiation, so that differentiating Equation (2) gives

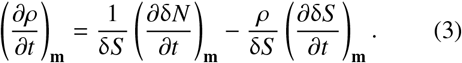

The first term on the right hand side of Equation (3) corresponds to changes in density induced by changes in the number of cells residing in δ*S*. The second term on the right hand side of Equation (3) describes changes in cell density due to local changes in the area of the portion of interface δ*S* during its evolution. In the first term, the number of cells may change due to proliferation, death, or net transport from the surrounding portions of the surface. The change in cell number due to proliferation and elimination can be expressed by (*P – A*)δ*N*(*t*) where *P* is the per capita proliferation rate, and *A* is the per capita cell elimination rate. The cell elimination rate may model cell death (apoptosis), detachment from the surface (for example anoikis), or embedment into the tissue. To describe the influence of tangential motion of the cells on cell density changes at position **r**_*s*_, we introduce the tangential flux of cells, **J**(**r**_*S*_, *t*). This cell flux is measured with respect to material points of the surface, which are themselves transported in space. It represents the number of cells crossing the boundary *C* of δ*S* per unit length per unit time (Figure 3a). The total number of cells leaving and entering δ*S* is thus calculated by the line integral of the flux of cells along *C*, where *C* is the curve surrounding δ*S*, with unit normal given by 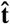 (Figure 3a). Therefore, the total rate of change of cell number in δ*S* is

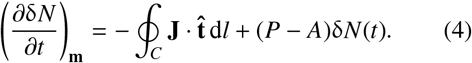

Since δ*S* is a small element of surface, the line integral in Equation (4) can be written in terms of the surface divergence of J, which can be formally defined as

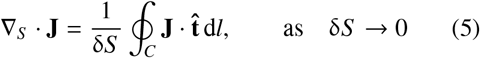

(Arnoldus, 2006). Thus, in the limit of an infinitesimally small area of the interface δ*S*, the change in density due to the change in number of cells in δ*S* in Equation (3) is given by

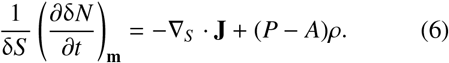

Equation (6) represents the fact that the surface divergence of the flux on a curved manifold is related to local changes in surface density (Arnoldus, 2006), much like, in the Euclidean space, the divergence of the flux is related to local changes in volumetric density.

To evaluate the second term of on the right hand side of Equation (3), we examine the rate at which δ*S* changes following the material trajectories of *S* (*t*). This depends on the local mean curvature,

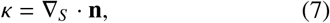

and is given by

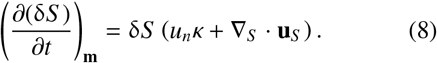

where **u**_*S*_ is the tangential component of the surface velocity and **u** (Figure 3). Equation (8) is derived using the equation for the change of a material area element over time from Batchelor (1967), see Appendix A for details. In our notation, *κ* is defined such that *κ* < 0 indicates concavity and *κ* > 0 indicates convexity.

Substituting Equations (6) and (8) into Equation (3), we find that the evolution of the surface density of cells following material trajectories of the interface is governed by

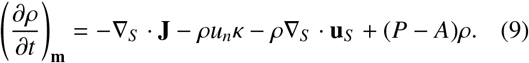

If cell migration includes advection and diffusion, the tangential flux of cells can be written as

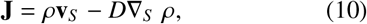

where **v**_*S*_ is the tangential velocity of the cells with respect to the surface and −*D∇_S_ ρ* corresponds to lateral diffusive flux along the curved interface where ∇_*S*_ is the surface gradient of *ρ*, that is the derivative of *ρ* on the manifold *S* (*t*) (Pressley, 2010). In this case, the evolution of the surface density of cells, Equation (9) becomes

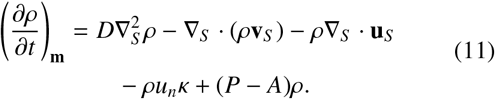

It is possible to determine the rate of change of cell density following other trajectories than the material points of the interface. The evolution equation for cell density takes a particularly convenient form when expressed following trajectories normal to the interface at each time (Figure 3). We can relate the derivatives of *ρ* along the normal and material trajectories by

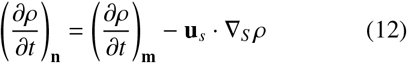

where (∂/∂*t*)_**n**_ represents the time derivative along the normal trajectories, that is trajectories perpendicular to the surface at all times (Wong et al., 1996). Substituting Equation (11) into Equation (12) gives

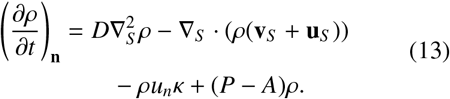

The first term on the right hand side of Equation (13) is the Laplace-Beltrami operator applied to the surface density of cells and describes the tangential diffusion of cells along the curved tissue surface (Berger, 2002). The second term describes the influence of tangential velocities of the cells **v**_*S*_ and of the tissue surface **u**_*S*_, respectively. The fourth term encapsulates the collective cell crowding or spreading effect of curvature, and the last term describes the gain or loss of cells from the group of tissue-synthesising cells. Equations (11) and (13) are general conservation equations for cells moving by advection and diffusion with respect to a surface which is itself moving and deforming. In Appendix B, we show that these equations are a generalisation of similar conservation equations of surface-bound quantities derived in the literature without tangential advection.

In our applications, for simplicity in the numerical solution, we will look at two dimensional problems where the interface is described by a one-dimensional tissue interface, that is, a curve in two-dimensional space. In these situations, Equation (13), can be written as

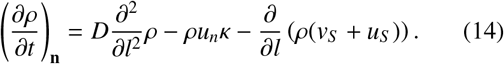

where ∂/∂*l* is the derivative with respect to the arc length of the surface, which is the one dimensional equivalent of the surface divergence and surface gradient (Redžić, 2001). In the applications presented in Section 3, we solve the coupled equations (1) and (14), where the tangential cell velocity is given various forms and the ensuing behaviour is analysed.

### 2.1. Numerical method

Solving Equations (1) and (14) requires solving a PDE on a moving boundary where the boundary motion is coupled with the PDE solution. To achieve this, we use an efficient hybrid computational method, the cell-based particle method (CBPM), developed in Leung and Zhao (2009), Leung et al. (2011) and Hon et al. (2014). In this method, the interface is represented by Lagrangian marker particles which are each associated with a grid cell of an underlying Eulerian grid with grid cell length Δ*x*. The grid is used to redistribute the particles along the moving interface to maintain quasi-uniform sampling. Furthermore, scalar quantities, such as cell density, can be associated directly with the marker particles (Leung and Zhao, 2009). This is an advantage over level-set like methods, which require additional scalar fields similar to the level-set function to represent surface-bound quantities (Alias and Buenzli, 2019). The interface is evolved over discretised timesteps Δ*t* by advecting the marker particles according to a velocity field. Local quadratic least squares interpolation of the interface and of the surface density of cells is then used to estimate the interface curvature and to evaluate spatial derivatives. The reader is referred to the Supplementary Information, Leung and Zhao (2009), Leung et al. (2011), Hon et al. (2014), and Hegarty-Cremer (2020) for more details.

## 3. Results

We now apply our mathematical model to cases of tissue growth where the inclusion of tangential cell advection allows us to model new biologically relevant situations. First, we validate the numerical method by solving simplified equations which test the two migration mechanisms of Equation (14), that is tangential cell advection and diffusion, as well as the crowding and spreading effect of curvature. These solutions are compared with analytic solutions. Then we model bone pore infilling and explore the generation of different orientations of collagen fibrils in infilled osteons, as illustrated in Figure 1b. We also model bone resorption, where osteoclasts tunnel through old bone tissue and investigate the influence of tangential cell velocity for the stability of travelling-wave-like resorption fronts observed during the resorption of cortical bone. Finally, we model the apical growth of root hairs and compare cell membrane trajectories and membrane curvature maps to experimental data (Shaw et al., 2000, Goriely, 2017).

### 3.1. Validation of the numerical method

To validate our implementation of the CBPM for solving Equation (14), we compare numerical simulations to analytical solutions in a simple setting where the governing equations are non-dimensionalised. To simplify the problem, the density is decoupled from the normal speed of the interface, that is we replace Equation (1) with *u_n_* = *c*, where *c* is a constant. We also set *D* = 0 and choose a circular initial interface with initial radius *R*_0_. In this case, the interface remains a circle at all times and it expands in the normal direction with radius *R*(*t*). We parameterise the circle using the arc length *l* and solve for *ρ* on the domain −π < *l* < π. The governing equations become

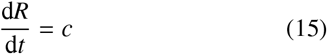

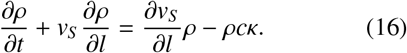

We assume an arbitrary initial cell density distribution *ρ*(*l*, 0) = *ρ*_0_(*l*) and an initial radius *R*(0) = *R*_0_, and impose periodic boundary conditions *ρ*(−π, *t*) = *ρ*(π, *t*). The solution for *R*(*t*) is

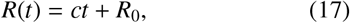

so that *κ*(*t*) = 1/(*ct* + *R*_0_). To test the advection term in Equation (14), we assume that cells are subject to the tangential cell velocity field *v_S_* = −*al* where *a* is constant. The governing equation for *ρ* becomes a quasilinear advection equation, which can be solved using the method of characteristics (Evans, 2010), giving

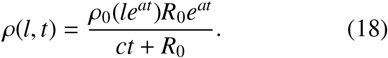

We test numerically both dilution of cells without advection, *a* = 0, and dilution of cells with advection, *a* ≠ 0. Figure 4 compares this analytical solution to the numerical solution obtained using the CBPM. In Figure 4, the initial condition for density is piecewise constant such that *ρ* = 0.5 when π/8 < |*l*| < 3π/8 and *ρ* = 0 elsewhere. There is excellent alignment between the analytic solution in Equation (18) and the one obtained by the CBPM both with and without tangential velocity. The small discrepancies are due to some degree of smoothing of the numerical solution, which originates from the local interpolation step of the CBPM. As expected, if the numerical discretisation is refined, the match improves. Convergence graphs can be found in the Supplementary Information, Figure S1.

**Figure 4:**
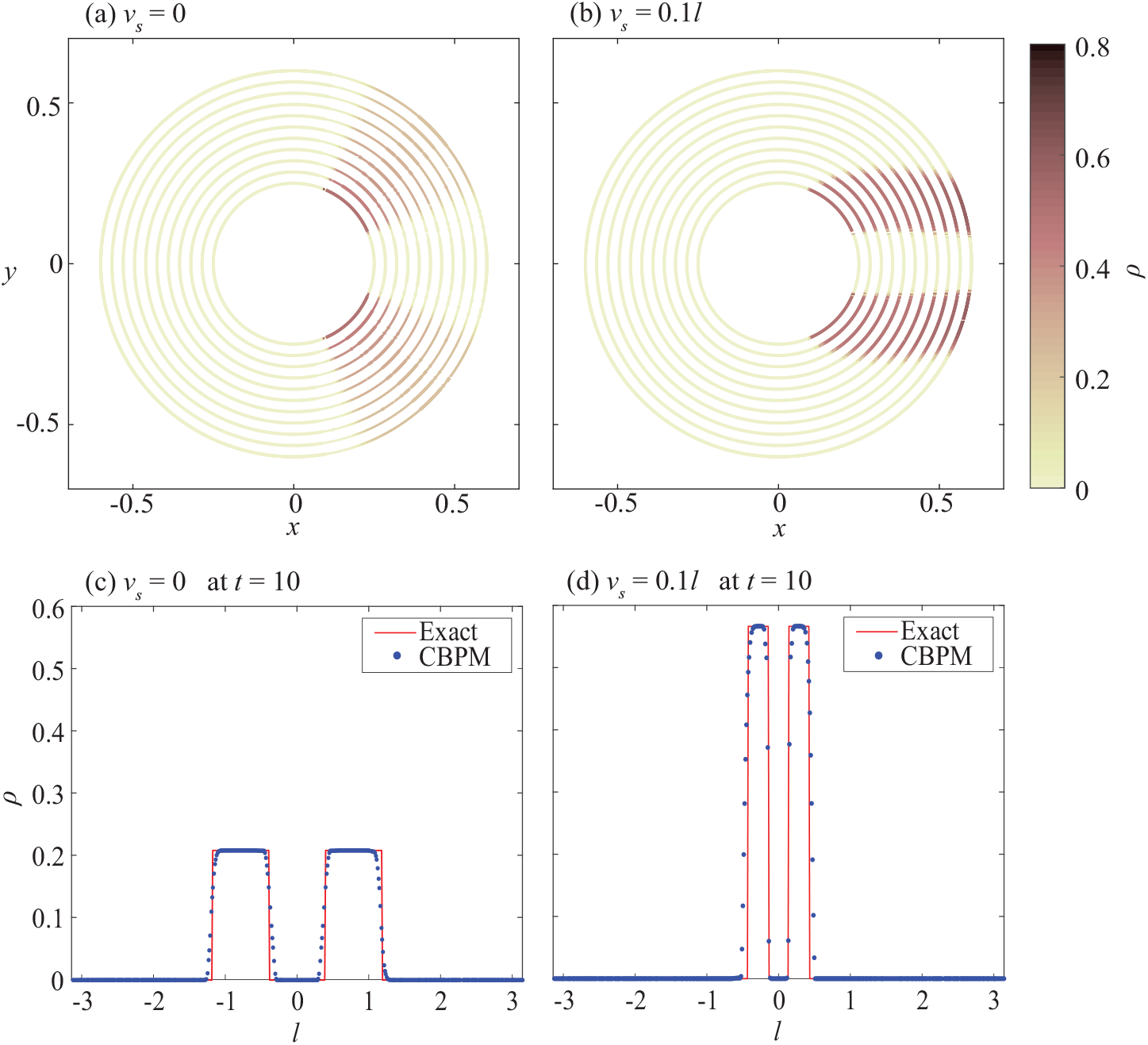
Expanding circle with *u_n_* = 0.035 and with or without tangential velocity: comparison between CBPM simulations and exact solutions. (a) and (b) Solution obtained using CBPM with *v_S_* = 0 and *v_S_* = 0.1*l*, respectively, with interface shown at regular time intervals (Δ*T* = 1). (c) and (d) Exact and CBPM solution density representation over arc length parameter at *t* = 10 with *v_S_* = 0 and *v_S_* = 0.1*l*, respectively. The discretisation used is Δ*x* = 0.01 and Δ*t* = 0.01.

To validate our implementation of the CBPM for problems that include diffusive transport, we solve the diffusion equation on a stationary circle using the CBPM. With a sinusoidal initial condition *ρ*_0_(*l*) = 0.5 + 0.5 sin(*l*) and periodic boundary conditions the analytic solution is given by

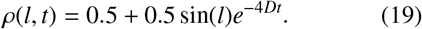

The results of the CBPM are compared with this solution at different times in Figure 5. Again, there is an excellent agreement between the solutions.

**Figure 5:**
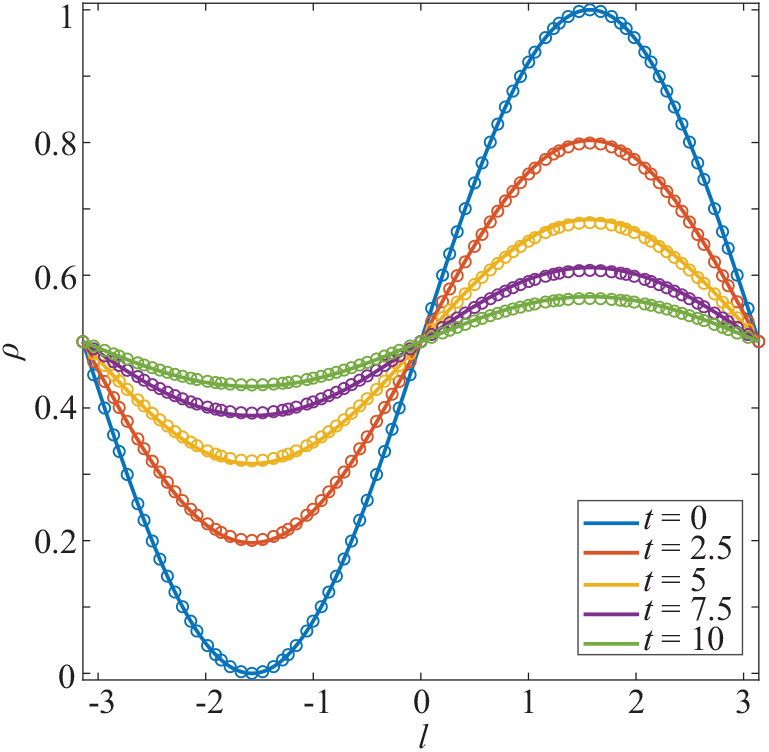
Analytic (solid line) and CBPM (points) solutions for diffusion around a stationary circle. The discretisation used is Δ*x* = 0.01 and Δ*t* = 0.01.

### 3.2. Circular bone pore infilling

We now consider the case of a circular bone pore being infilled by a population of osteoblasts distributed uniformly along the pore’s perimeter. This can be thought of as the infilling of a cortical bone osteon seen in a transverse cross section. New bone tissue is gradually produced such that the initial interface is moving inwards while retaining a circular shape. As infilling proceeds, the density increases as a result of the systematic effect of curvature (Buenzli, 2014, 2016). We examine three cases of tangential cell velocity: no tangential velocity, constant tangential cell velocity, and time-dependent tangential cell velocity such that cells reverse their motion with respect to the interface at specific times (Figure 6). By rotation symmetry, in these simulations, the density remains uniform at all times, but it is time dependent due to the shrinkage of the bone surface area as infilling proceeds. The model parameters in Figure 6 are based on experimental values, see Alias and Buenzli (2018) for more details.

**Figure 6:**
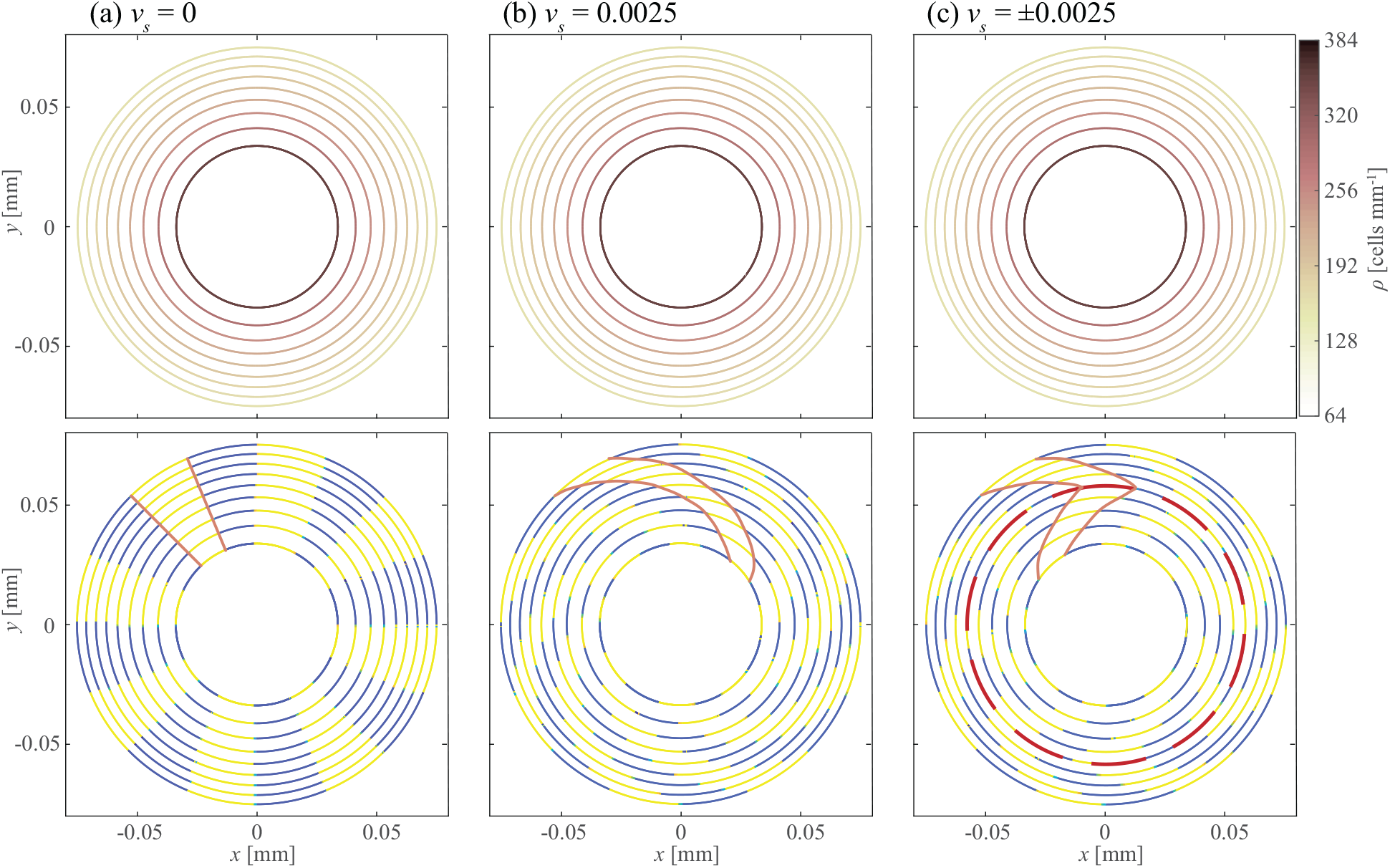
Circular pore infilling results with cell secretion rate *k* = 7.8125e 06 mm/day and with varying tangential cell velocity. In each figure the initial interface is the outermost ring and the interface is shown at regular time intervals (Δ*T* = 3 days). The top figures show density and interface position while the bottom figures show cell trajectory tracking and interface position with a single cell trajectory annotated in orange. (a) Infilling circle without tangential velocity. (b) Infilling circle with tangential velocity *v_s_* = 0.0025 mm/day. (c) Infilling circle with tangential velocity *v_s_* = 0.0025 mm/day when *t* < 12.5 days and *v_s_* = −0.0025 mm/day when *t* ≥ 12.5 days. The location of the change of direction is emphasised in a red dashed circle. The discretisation used is Δ*x* = 0.001 mm, Δ*t* = 0.075 days.

The evolution of density and interface position is the same across the three cases (Figures 6a–c). However, the cell trajectories in space are distinct, and this creates different tissue material properties (Figures 6d–f). To visualise cell trajectories in Figures 6d–f, cells are stained either in blue or in yellow. This is achieved in the CBPM by assigning a new scalar property to each marker particle, which is simply advected along the cell trajectories. In Figure 6d, cells have no tangential motion hence their trajectories are moving along straight radial lines. However, in Figures 6e and 6f, the cells move tangentially to the surface, thus their trajectories spiral inwards. The results in Figure 6f illustrate how one may explain a change in anisotropic tissue material properties. As collagen fibrils secreted by osteoblasts may be weaved according to the directionality induced by cell migration, the change in cell trajectory orientation could be used to describe the change in collagen fibre orientation in lamellar bone and the consequent plywood structure (as illustrated in Figure 1b).

### 3.3. Bone resorption in basic multicellular units

We now examine the resorption phase of a bone remodelling event as another example where the tangential velocity of cells may be important for the evolution of the tissue interface. In bone resorption, bone tissue is removed by osteoclasts attached to the bone surface. The resorption of bone matrix by osteoclasts creates a cavity which maintains consistent cellular organisation and shape at the resorption front (Figure 1c) (Jaworski and Hooper, 1980, Ryser et al., 2009, Buenzli, 2010, 2011, 2014, Buenzli et al., 2012, Lassen et al., 2017). We apply our tissue growth model to this situation to show that to maintain this stable travelling resorption front, directed tangential osteoclast motion is required (Figure 1c).

Recent works have suggested that osteoclasts at the front of basic multicellular units may remain at this position for a long period of time (Lassen et al., 2017), unlike previous suggestions that osteoclasts move down the cavity walls (Burger et al., 2003, Buenzli et al., 2012). We show here, based on simple numerical simulations, that a stable resorption front requires cell guidance mechanisms to steer osteoclasts back toward the tip of the cavity (Figure 1c). Without such directed motion, the cavity rapidly expands out and osteoclasts move away from each other (Figure 7a). Figures 7b and 7c show numerical simulations where two different types of signals are used to steer osteoclasts back toward the tip of the cavity. The first signal modelled can be thought of as haptotaxis, which is a cell guidance mechanism in response to adhesion gradient on the substrate generated by cell binding to substrate molecules (Davies, 2013). The second is chemotaxis, which describes cell guidance through a chemical gradient (Murray, 2002). These signals inducing cell tangential velocity could originate from mechanical signals, believed to be important in guiding bone remodelling processes in bone. Around the tip of the resorption cavity, mechanical stresses are increased (Smit and Burger, 2000, Ruimerman et al., 2005). Osteocytes, which form a network within bone matrix are able to sense mechanical deformation and transduce these mechanical variables into molecular signals. These molecular signals may then diffuse through bone matrix and in the resorption cavity, where they are sensed by the osteoclasts as a chemotactic signal. Alternatively, mechanical gradients along the resorption cavity walls may be felt directly by osteoclasts as a haptotacic signal.

**Figure 7:**
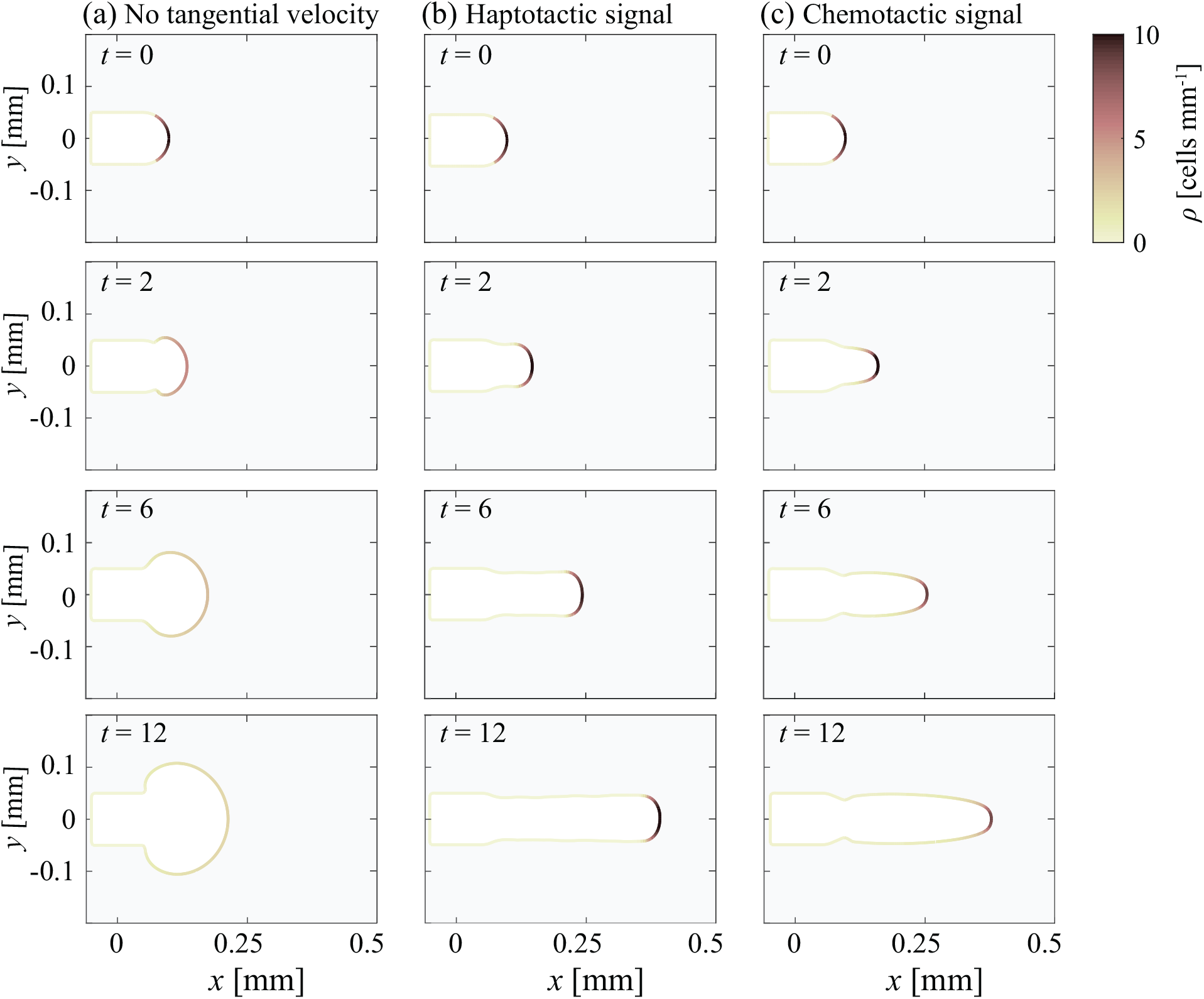
Bone resorption results with different forms of tangential cell advection. The time is shown in days and the spatial unit is mm. The resorption rate is 0.025 mm/day (*k* = −0.025). (a) Resorption front behaviour with no tangential velocity pulling cells. (b) Haptotactic signal: arc length dependent tangential velocity. The proportionality constant between the arc length distance and the tangential velocity is *a* = 0.6. (c) Chemotactic signal: tangential velocity determined by the projection of an external gradient field on the interface. The discretisation used is Δ*x* = 0.00375, Δ*t* = 0.02.

Osteoclasts work in close contact with other cells lining the cavity walls, called reversal cells, which may provide haptotactic signals such as receptor activator of nuclear factor kappa-B ligand (RANKL) (Martin et al., 2004, Lassen et al., 2017). Here we assume the haptotactic signal induces a tangential velocity to the osteoclasts, *v_s_* = *a l*, where *l* is the arc length measured along the cavity wall from the tip and *a* is a positive constant. This is similar to Section 3.2 where *l* > 0 on the upper part of the cavity and *l* < 0 on the lower part of the cavity. Using this form of tangential cell velocity, it can be seen from Figure 7b, that a stable resorption front is formed. Between *t* = 0 and approximately *t* = 3 days, there is a transient period, where the shape of the resorption front evolves until a balance between the advection-induced crowding and curvature-induced spreading of the osteoclasts is achieved. After this transient, the cell density profile and the cavity front shape is maintained as it progresses through the bone tissue.

Alternatively, we model chemotaxis by projecting a velocity gradient field, such as one created by a gradient of chemical concentration −*b*∇*C*, onto the cavity surface and taking this projection as the tangential velocity,

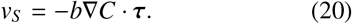

Indeed, active osteoclasts remain bound to the interface, therefore they can only explore the tangential component of the chemical gradient field. This gradient could be due to signalling molecules derived from mechanically-stimulated osteocytes embedded in bone matrix, that steer osteoclasts toward specific areas of bone needing repair (Turner et al., 1994, Marotti, 2000, Ryser et al., 2009, Lerebous and Buenzli, 2016), such as high mobility group box protein 1 (HMGB1) (Yang et al., 2008) and colony-stimulating factor 1 (CSF-1) (Harris et al., 2012), or it may be due to other chemotactic molecules from the bone microenvironment, such as monocyte chemoattractant protein-1 (MCP-1/CCL2) (Wu et al., 2013), and the chemorepulsing sphingosine-1-phosphate (S1P) (Ishii et al., 2010). For the results presented here, we simply take −*b*∇*C* = [0, −2.5 sgn(*y*)*y*^2^], which is a velocity field in the *y* direction with streamlines pointing towards the centerline of the cavity. Figure 7c shows that, similarly to the haptotaxis results, stable resorption front behaviour is obtained after an adjustment period between *t* = 0 and *t* = 3 days.

Both forms of cell guidance signal result in stable resorption fronts, but they lead to different resorption cavity shapes, indicating that the type of signal is also important for the resorption front. The chemotactic signal results in a wider distribution of osteoclasts around the resorption front compared to the haptotactic signal, which results in a high concentration of cells on a narrow portion of the interface. Due to coupling, these differences in cell densities are reflected in the shape of the resorption fronts. However, the speed of these resorption fronts is comparable, with the haptotactic signal canal reaching *x* ≈ 0.345 mm at *t* = 12 days and the chemotactic signal canal reaching *x* ≈ 0.34 mm at *t* = 12 days. These speeds align well with expected speeds of resorption cavities (30-40 μm/day) (Jaworski et al., 1981, Lassen et al., 2017). A combination of both signals is also possible, and leads to similar conclusions (Supplementary Information, Section S2). The effect of changing parameter values for the haptotactic and chemotactic signals is also shown in the Supplementary Information (Figure S2).

The diameter of the simulated resorption cavities falls within the values stated in the analysis of exerimental resorption cavities by Lassen et al. (2017). In our simulations, the ‘Level 1’ canal diameters (25 μm from the front) are 58.8 μm for the chemotaxis and 87.7 μm for the haptotaxis, and the ‘Level 2’ canal diameters (325 μm from the front) are around 100 μm for both chemotaxis and haptotaxis. The range of diameters found in Lassen et al. (2017) for Level 1 is 30–180 μm with the mean being 80 μm and for Level 2 it is 110–390 μm with the mean being 200 μm.

### 3.4. Root hair growth

We now apply our model to the apical growth of root hairs. Root hairs have a single tip-growing cell which concentrates secretory vesicles to the tip of the cell (Miller et al., 1997). To apply our model to this situation, we take the underlying surface to be the root hair cell membrane, and the surface density *ρ* to represent the density of secretory vesicles near the root hair cell membrane (Shaw et al., 2000). We assume that the cell membrane only has a normal velocity, as described by Equation 1 and as suggested by experimental observations (Figure 8c), and that the secretory vesicles have a tangential velocity determined by a haptotatic signal (*v_s_* = *al*). This tangential velocity allows the position of the vesicles to be maintained near the tip of the cell (Shaw et al., 2000).

**Figure 8:**
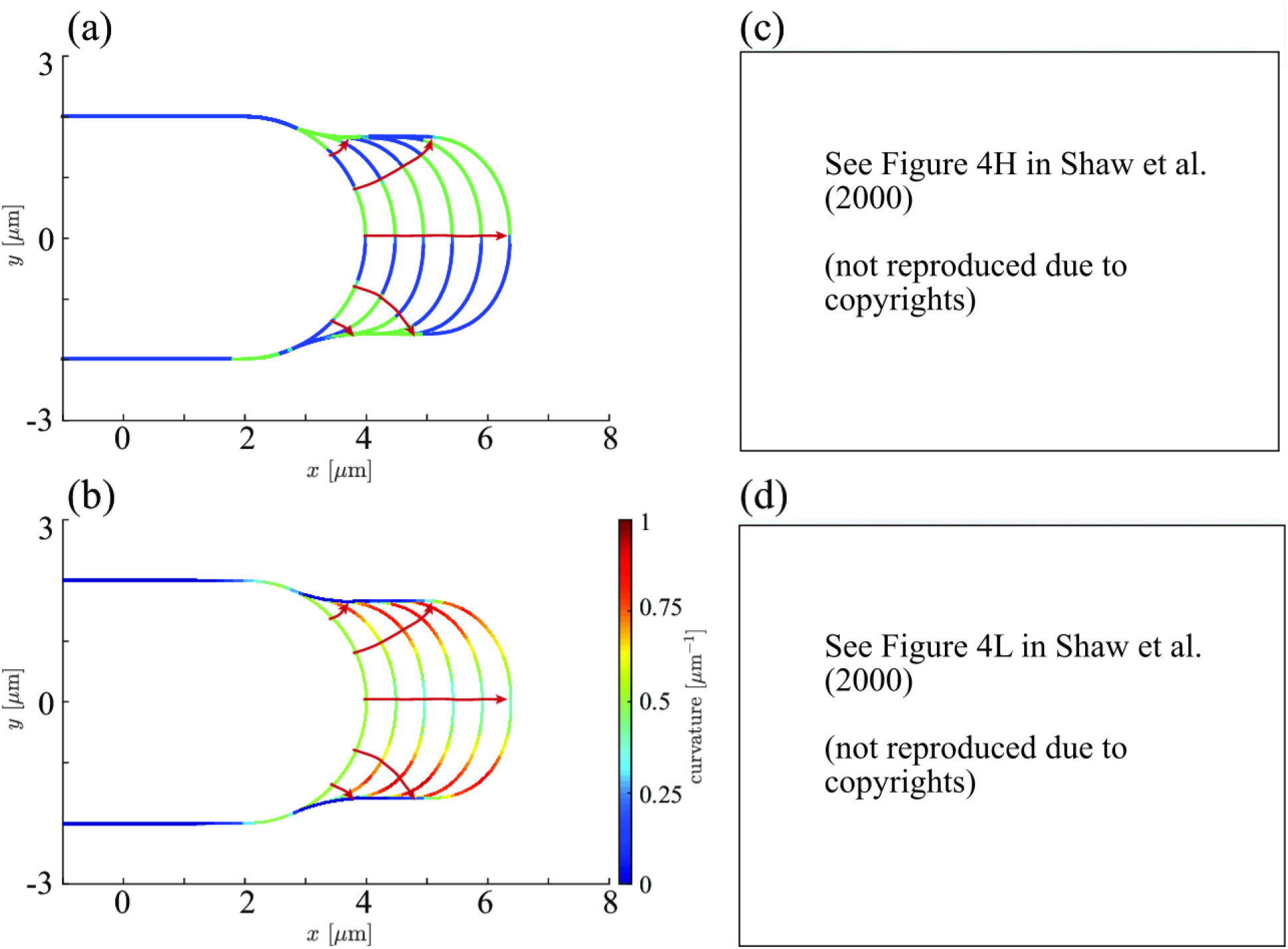
Apical growth of root hair cells. (a) The numerically simulated root hair cell membrane is shown every 5 minute intervals, assuming a secretory rate of 0.1μm/minute. Material points of the cell membrane are stained in blue or green and their trajectories are highlighted in red. (b) Curvature map along the numerically simulated cell membranes. (c) Experimental material point tracking results (green) for a 36 minute tracking experiment (reproduced with permissions from Shaw et al. (2000)). (d) Curvature map versus arc distance from the root hair centre over time (reproduced with permissions from Shaw et al. (2000)).

By tracking the material points of the root hair cell membrane in time in our model, we see very similar trajectories to those observed in the growing root hairs analysed in Shaw et al. (2000), where microspheres adhering to the cell membrane were used to track the evolution of its material points (Figures 8a and c). Without tangential velocity, the secretory vesicles would also follow these trajectories and the cell would not grow its tip only in the longitudinal direction. Furthermore, the curvature of the membrane around the growing tip reflects the curvature map measured in growing root hairs (Shaw et al., 2000), where highest curvature is found a small distance away from the tip centre (Figures 8b and d). This example application shows that the addition of tangential velocity to the tissue growth model allows for multiple different new applications of tissue growth to be explored.

## 4. Discussion and conclusion

Tangential cell motion generated by cell guidance mechanisms is important in several situations of tissue growth, such as growth occurring at an angle with respect to the tissue surface, and the generation of anisotropic tissue properties. We have developed a new mathematical model for tissue growth under collective curvature control to incorporate such directed cell guidance mechanisms by including tangential cell motion. The model is derived from conservation principles applied to the surface density of tissue-synthesising cells. This derivation results in a PDE for cell density on a moving boundary, which is coupled with the boundary motion. The governing equations are expressed in covariant form, that is, they are independent of a choice of surface parameterisation and coordinate system. We solve the model numerically using a hybrid front-tracking computational method, the CBPM, and find good agreement with analytic solutions.

Experimentally, the interaction between curvature control of tissue growth and directed cell motion is difficult to investigate, due to the challenge of controlling evolving tissue geometries. Crowding and spreading effects on rates of tissue progression are a consequence of space constraints that may be masked by cell behavioural influences in experiments. The development of mathematical models that account for such collective effects can help disentangle geometric and cell behavioural influences of tissue growth (Cai et al., 2007, Alias and Buenzli, 2018, Buenzli et al., 2020). The example of bone tissue resorption developed in this paper (Figure 7) illustrates the importance of taking into consideration both the mechanistic influence of curvature on osteoclast density, and the tangential motion of osteoclasts with respect to the bone interface. Without accounting for the mechanistic influence of curvature, the presence of a driving force steering osteoclasts toward the centerline of the resorption cavity would not be highlighted. Without tangential motion of osteoclasts at the tip of bone resorption cavities, our results suggest that stable cavity shapes are not possible.

Our mathematical model describes the joint evolution of the tissue interface and tissue-synthesising cell density. The example of bone pore infilling in Figure 6 illustrates that directed motion of cells can generate anisotropies in tissue material properties. While we did not model tissue generation explicitly, our model may be coupled with more detailed tissue generation mechanisms that include creation and destruction of tissue material at moving interfaces, as well as tissue maturation mechanisms, based on bulk and surface mass balance (Cumming et al., 2010, Buenzli, 2015, 2016). Our model thus provides a basis for further explorations into the relationship between the spatial organisation of anisotropic tissue material properties, and the dynamics of their creation. Biological experimental data often takes the form of tissue samples or biopsies representing single snapshots in time of the state of the tissue. This type of data contains detailed spatial information about the organisation of a tissue, but it does not offer a detailed picture of its time evolution. The provision of mathematical links between features recorded in the state of a tissue and the dynamics of its formation may allow us to deduce how a tissue has been produced given an analysis of its material properties. In bone tissues, for example, several features of bone formation are recorded, such as osteocyte density (Buenzli, 2015), mineral density (Buenzli, 2016, Lerebours et al., 2020), and tetracycline labels and lamellae, which provide information about past location of the bone interface (Martin et al., 2004, Buenzli, 2014, Andreasen et al., 2018). This type of information is used in bioarcheology to estimate archaeological age and activity (Buckberry and Chamberlain, 2002, Maggiano et al., 2008, Mays, 2010). An analysis of lamellae patterns in bone cross sections could provide more information about osteoblast behaviour, and provide more insights in cases of irregular bone formation patterns such as drifting osteons (Robling and Stout, 1999, Maggiano, 2012) and bone disorders.

Discretising PDEs on moving boundaries is a challenging problem of applied mathematics. In this paper, we restricted our model to two dimensional applications for simplicity. Clearly, applications of our model to three-dimensional tissue growth are of interest (Figure 1) (Guyot et al., 2014, Goriely, 2017, Ambrosi et al., 2019, Ehrig et al., 2019). Sophisticated techniques have been developed to simulate the evolution of interfaces in three-dimensional complex systems (Sethian, 1999, Tryggvason et al., 2001, Glimm et al., 2001, Shin and Juric, 2002, Osher and Fedkiw, 2003, Du et al., 2006, Leung and Zhao, 2009, Hon et al., 2014). While the level-let-like method developed in (Alias and Buenzli, 2019) for curvature-controlled tissue growth may be suitably adapted to include tangential cell velocity, the CBPM of Hon et al. (2014) used in this work is also applicable to three-dimensional interfaces.

## Acknowledgements

This work is supported by the Australian Research Council (DP180101797, DP200100177), the Centre for Biomedical Technologies, Queensland University of Technology (QUT), and the Institute of Health and Biomedical Innovation (IHBI), QUT, as well as the VELUX foundation (Project no. 25723). We thank the three anonymous referees for their helpful comments.

## Appendix A. Evolution of local surface area

We start with the equation for the rate of change of a vector area element (δ**S**= **n**δ*S*) of a material surface from Batchelor (1967),

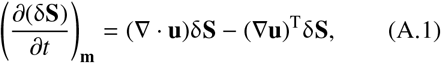

where ∇**u** is the Jacobian matrix of **u**. Following Stone (1990), we take the inner product with **n**, to obtain an expression for the change in local surface area, δ*S*, over time,

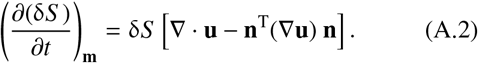

The right hand side of Equation (A.2) corresponds to subtracting to the total divergence of **u**, that is, to the trace of the Jacobian matrix of **u**, the normal component of the trace, **n**^T^(∇**u**) **n**= **n**^T^(∇**u**)^T^ **n**. This gives the surface divergence operator of **u**, so that

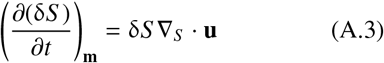

Decomposing **u** into its tangential and normal components, **u** = *u_n_***n** + **u**_*S*_, one gets

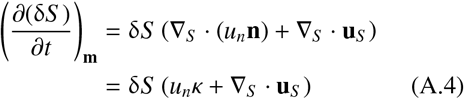

where the second equality in Eq (A.4) uses the fact that the surface divergence of the unit normal vector is the mean curvature of the surface, *κ* = ∇_*S*_ · **n**(Goldman, 2005), and that the surface gradient is perpendicular to **n**, so that **n** · ∇_*S*_ *u*_*n*_ = 0.

## Appendix B. Comparison with the literature

In multiphase physico-chemical systems, similar evolution equations to Equation (11) are derived for the surface transport of surfactants at the interface between two phases (Stone, 1990, Wong et al., 1996, Xu and Zhao, 2003). A difference between such physical systems and the biological systems we are modelling is the coupling between the surface velocities and the cell density via Equation (1). Cell density affects interface evolution, whereas in multiphase physico-chemical systems, surface evolution is usually assumed to be independent of surfactant density. Furthermore physico-chemical system models do not consider the tangential velocity of a surfactant with respect to the surface.

In Stone (1990), surfactant mass balance equations are derived, however the nature of the time derivative of surfactant density is unclear (Wong et al., 1996). Time derivatives in Stone (1990) implicitly represent changes following paths normal to the interface. The surfactant mass balance results obtained in Wong et al. (1996) make the nature of the time derivative explicit by being derived using an explicit parameterisation of the interface. The parameterisation is general in the sense that the coordinate system is not necessarily bound to the material points of the interface. If we set **v**_*S*_ = 0, *A* = *P* = 0 in Equation (13), we fall back on Equation (7) from Stone (1990) following normal trajectories, and Equation (5b) from Wong et al. (1996) equation if the timelines of their parameterisation are taken to be following the normal trajectories of the surface.

Neither the equations in Wong et al. (1996) nor Stone (1990) include coupling between the interface speed and density of cells nor tangential velocity. The derivation in Alias and Buenzli (2017) includes coupling between cell density and interface speed, but the cells have no tangential advection, that is, their only lateral motion is diffusive. To compare our model with that in Alias and Buenzli (2017), the cell velocities in Equation (13) must be chosen such that the cells move along the normal trajectories of the interface. Therefore, if we set *v_S_* = −*u_S_*, the governing equations agree.

## Supplementary Information

### S1. Numerical Discretisation

We provide more detail on the numerical method used to solve our mathematical model, the cell-based particle method (CBPM). As well as the advantages of the CBPM discussed in the main text, the CBPM also allows for efficient detection and implementation of topological changes during fusion or fragmentation of the interface without requiring information about the connectivity of the marker particles. This method is of 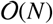 in computational load, where *N* is the number of marker particles. This is to be contrasted with standard level set methods on *N × N* grids, which have 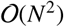 computational load, or 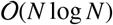 for local level set methods (Sethian, 1999). The CBPM method for solving PDEs on moving boundaries gives approximately conservative solutions (Leung and Zhao, 2009).

The CBPM algorithm is comprised of four main steps: initialisation, movement, resampling, and activation or deactivation. The initialisation stage declares the set of active marker particles which are used to track the interface, its geometry, and any associated scalar quantities. Initialisation requires an explicit parameterisation of the initial surface, *γ*(*m*), where *m* is a parameterisation variable for the one-dimensional surface *γ* in two dimensional space. The movement stage requires a velocity field which is used to advect the marker particles, and thus to evolve to position of the interface. This velocity field can be space and time dependent, and may be determined by external processes, or be coupled with the evolution of intrinsic properties of the system. In our case, it is intrinsic and implicitly time dependent since it depends on the surface density of cells. To solve the partial different equation (PDE) for a scalar quantity residing on the moving interface, each marker particle is supplemented with a scalar value, which is evolved according to the particular PDE after the motion step of the marker particles.

The resampling stage assures a quasi-uniform sampling of the interface through local interpolation. The local interpolation of the interface is expressed in a local coordinate system aligned with the local surface unit normal and calculated using quadratic least squares. The interpolation is used to update the unit normals, curvature, and any other local surface properties of interest, as well as to resample the active marker particles and calculate spatial derivatives in Equation (13). Finally, the activation and deactivation stage deactivates marker particles associated with underlying grid cells which no longer contain part of the interface, and activates marker particles associated with underlying grid cells into which a portion of the interface has now moved. The activation and deactivation stage also detects changes due to topological changes of the interface incldugin collision, fusion, or fragmentation. For more details on the algorithm see to Leung and Zhao (2009), Leung et al. (2011), and Hon et al. (2014). Below we describe how the method is applied to our problem.

To solve Equations (1) and (13), we first need to choose an advective velocity field in the two dimensional space. Several choices are possible, including paths normal to the interface at all times, material points of the interface, and cell trajectories. Since cells may carry intrinsic information and it is expected that the numerical resolution of cell density changes will be more accurate along these trajectories, we choose to move the marker particles along cell trajectories. We thus define the intrinsic velocity field that governs the evolution of the surface by

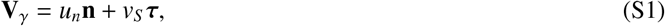

and we use a forward Euler scheme to evolve the positions of the marker particles in time for simplicity. More advanced time stepping schemes can be devised (Leung and Zhao, 2009, Leung et al., 2011, Hon et al., 2014), but in practice, moving boundary problems are more sensitive to spatial discretisation accuracy than time discretisation accuracy (Osher and Fedkiw, 2003). Equation (13) is solved after interface motion using operator splitting with forward Euler, with the first step solving for curvature control,

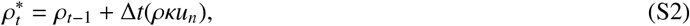

where the * indicates an intermediary step in the solution for *ρ*_*t*_ and Δ*t* is the time step. Given there is a local interpolation for both the surface and the density values, denoted by

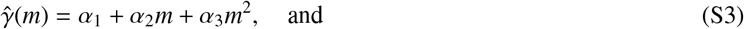

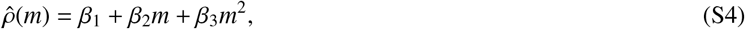

respectively, we can then calculate the diffusion term of Equation (13) by calculating the Laplace Beltrami operator

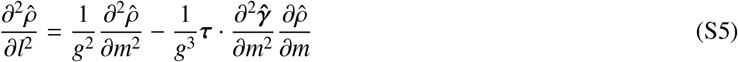

directly from the second order interpolation, 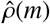. In Equation (S5), *g* is the surface metric, 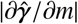. Similarly to the Laplace-Beltrami operator, the final term of Equation (13) can be calculated using the local interpolation. The derivative of *v_S_* with respect to *l* can be calculated either explicitly or via interpolation depending on the form of *v_S_*. The forward Euler method is then used to step the diffusion and advection operators forward in time,

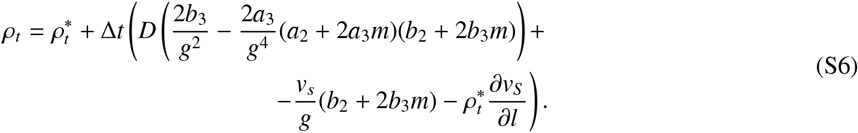

**Figure S1:**
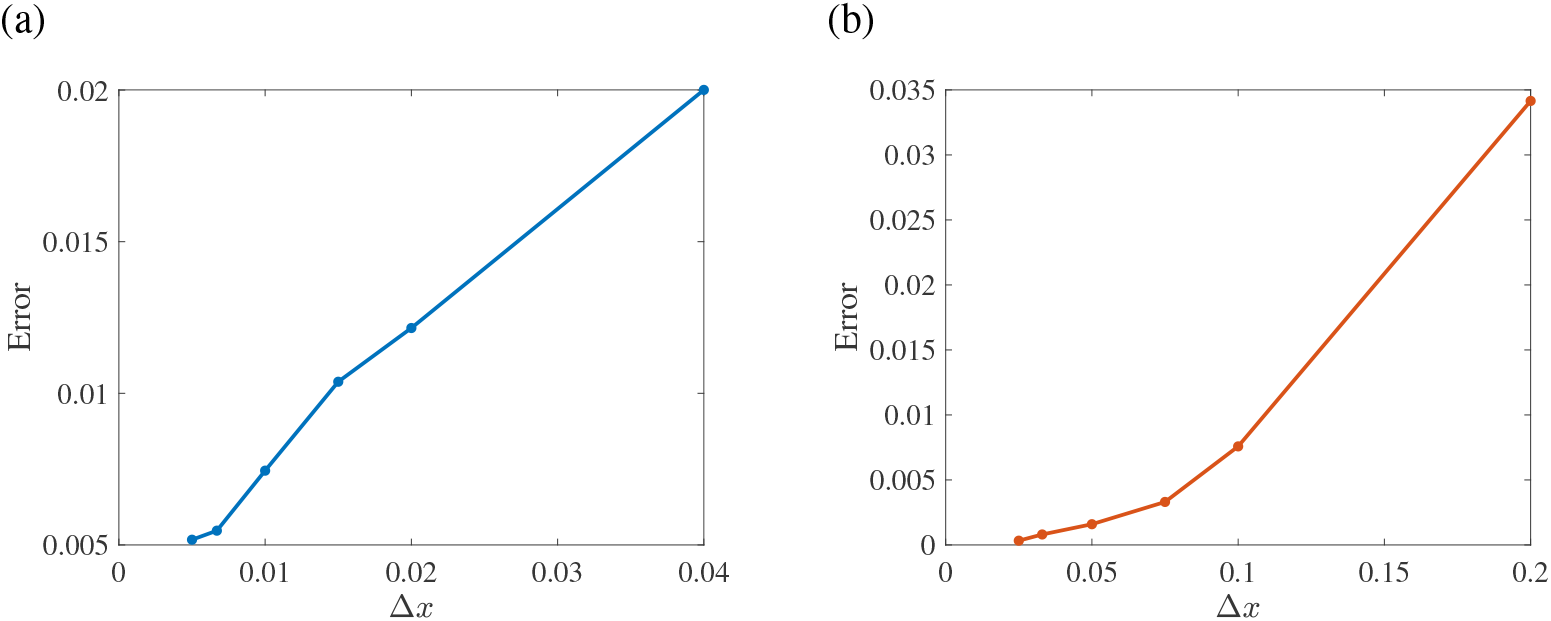
Discretisation convergence graphs. (a) Comparison of average absolute error of the advection-only problem (Equation 18). The time step is changed with the spatial step, maintaining the relation Δ*t* = Δ*x*. (b) Comparison of average absolute error of the diffusion-only problem (Equation 19). The time step is changed with the spatial step, maintaining the relation Δ*t* = Δ*x*^2^/(2*D*).

This concludes the calculation of *ρ* between timesteps.

We also include convergence graphs in Figure S1, where we examine the error of the final time solution compared to the analytic solutions presented in Figures 4 and 5. The error is calculated by comparing the densities at *t* = 10 along the arc length of the surface. For each discretisation, the absolute error is calculated for every marker particle, then the average of these absolute errors is found. The figures show that indeed the error is reducing as the discretisation is refined, and the spatial discretisations chosen for the results presented above are justified.

**Figure S2:**
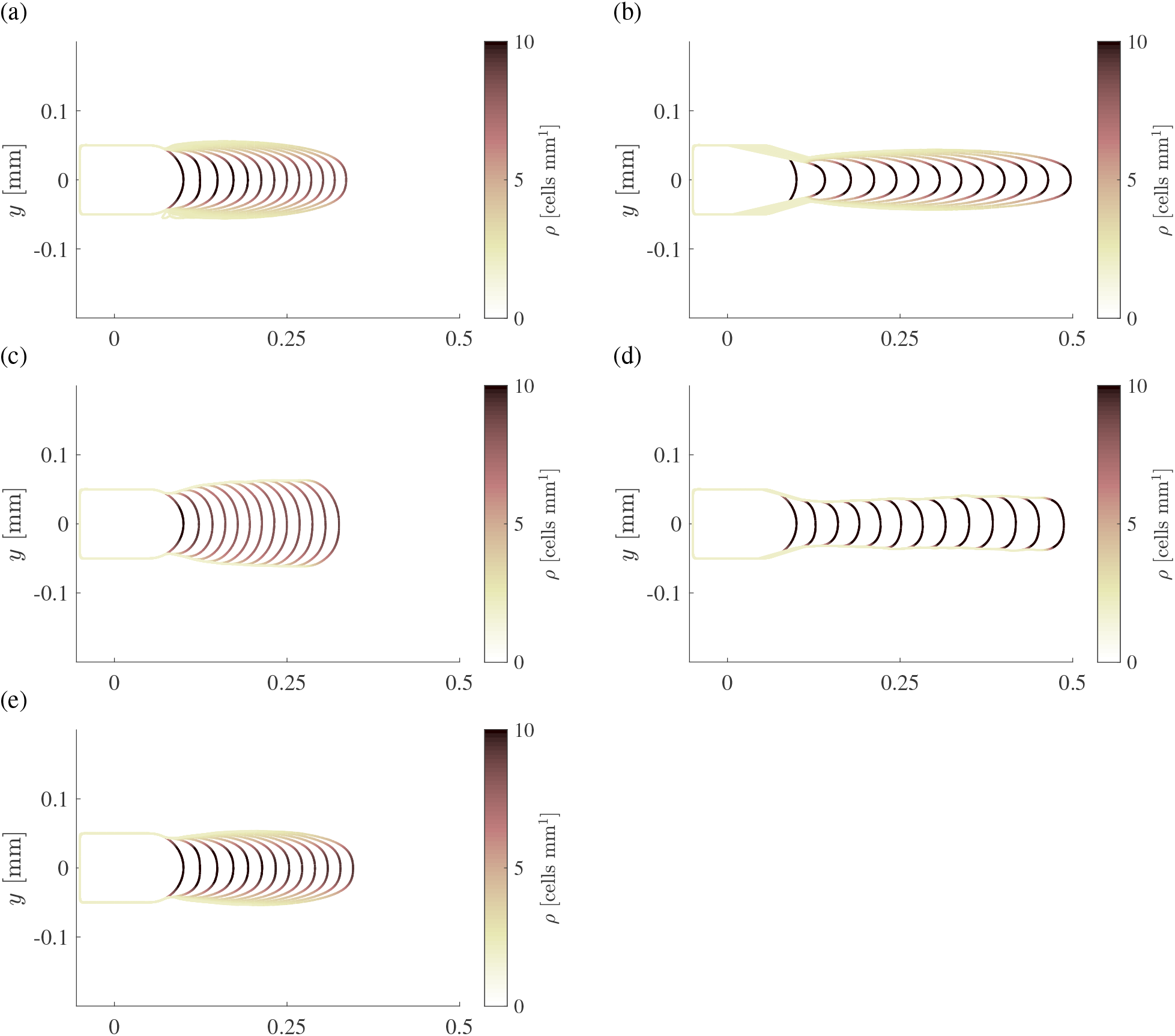
Sensitivity analysis of resorption behaviour to proportionality constants. (a) and (b) Chemotactic signals with proportionality constants of *b* = 1.25 and *b* = 5 respectively. (c) and (d) Haptotactic signals with proportionality constants of *a* = 0.3 and *a* = 1 respectively. (e) Combination of chemotactic and haptotacitc signals with *b* = 1 and *a* = 0.1.

### S2. Effect of resorption constants

Here we present results which show the sensitivity of the resorption behaviour discussed in Section 3.3 to the strength of the cell signals. We present results for different values of *a*, the proportionality constant of the haptotactic signal, and different values of *b*, the proportionality constant of the chemotactic signal. We also present the result if both signals are superposed linearly. For both signals, we see that increasing the constant of proportionality increases the strength of the signal and causes a narrower front which travels faster through the bone (Figure S2b and c). Weakening the signal has the opposite effect (Figure S2a and c). Superposing the two types of signals exhibits elements of the behaviours of each. There is a flatter front of the cutting cone, which can be seen in the haptotactic results, but there is also a widening of the channel about 100μm from the front which is characteristic of the chemotactic results. Therefore, through the combination of these two signals, many different resorption behaviours can be modelled.

